# Mitigating Family Effects in RNA Secondary-Structure Prediction with Latent-Space Continual Learning

**DOI:** 10.64898/2026.04.29.721709

**Authors:** Wissal Mokeddem, Giulia Pedrielli, Teresa Wu

## Abstract

Accurate RNA secondary-structure prediction remains difficult despite decades of thermodynamics-based algorithms and the advent of deep-learning architectures (convolutional networks, Transformers, diffusion models). In fact, the datasets that pair RNA sequences with secondary-structure labels are often low-quality, noisy, and family-imbalanced, which limits out-of-distribution generalization and exacerbates catastrophic forgetting when new data regimes are introduced. We propose a continual-learning approach based on Lifelong Bayesian Optimization (LBO), RNAFOLBO, that treats each class of RNAs obtained from latent-space clustering as a sequential task and jointly orchestrates training and hyperparameter selection of heterogeneous models (UFold, RNA-FM, RNADiffFold), while preserving prior knowledge. Concretely, we apply LBO to 15 clusters obtained by clustering RNAStrAlign in the latent space of RNAGenesis, a model specialized in contextual representation learning and latent-space structuring, achieving a mean *F* 1 per cluster of 0.931 (with a range of 0.177). These results surpass the strongest one-shot baseline and mitigate forgetting without full retraining. The gains persist as additional clusters are introduced. Overall, RNAFOLBO delivers higher and more stable performance and practical scalability for integrating new RNA clusters or families, enabling more robust and transferable RNA secondary-structure prediction.

## Introduction

Ribonucleic acid (RNA) molecules are essential macro-molecules that perform diverse biological functions in living organisms (1). From genetic information transfer and gene regulation to catalysis by ribozymes and metabolite sensing by riboswitches, RNA has a role to play (2, 3). Therefore, a deeper understanding of RNA structure is a major priority for many fields such as structural biology, synthetic biology, virology, drug discovery, and molecular diagnostics.

Primary, secondary, and tertiary structures each matter: the primary structure (the nucleotide sequence) determines local pairing propensities and constrains the space of feasible folds. The research concerned with primary structure looks into sequence representation learning (RNA language models / embeddings). The goal is to learn contextual, transferable features from unlabeled RNA sequences (motifs, constraints, long-range dependencies) that support structure/function pre-diction. RNA-FM is a widely cited example of an RNA-specific foundation model used for downstream tasks including secondary-structure prediction (4). Another field that is relevant to primary structure is RNA design / inverse folding (primary structure generation conditioned on structure or function). Here the task is to produce sequences that realize a target secondary structure or function. A well-known demonstration that community-driven and algorithmic methods can derive strong design principles is the Eterna “massive open laboratory” work (5). This literature is relevant because (i) it emphasizes that sequence–structure mapping is many-to-one and (ii) it motivates methods that reason over distributions and regimes rather than a single deterministic mapping. Importantly, sequence-level models and embeddings provide a natural space for clustering RNAs into regimes/tasks. Under this view approaches such as continual learning represent a principled way to update predictors as new regimes appear.

The secondary structure specifies the set of intramolecular base pairs (often represented as a contact matrix or dot–bracket string), capturing the dominant energetic contributions and many conserved functional motifs.

Unlike secondary structure, tertiary folding is strongly shaped by context-dependent stacking, electrostatics of the phosphate backbone, metal ions, and complex loop–loop/receptor interactions, making the search space highly rugged and the data comparatively scarce. Community-wide blind challenges such as RNA-Puzzles have therefore played a central role in objectively assessing progress and exposing persistent bottlenecks in modeling noncanonical motifs, coaxial stacking, and global topology (6). On the methods side, leading approaches include fragment/motif assembly and physics-inspired refinement pipelines—exemplified by Rosetta FARFAR2, which consolidates puzzle-driven innovations and achieves native-like folds on many targets while still struggling on larger RNAs due to sampling and scoring limitations (7). More recently, deep-learning models have expanded toward RNA and protein–nucleic-acid complexes, with RoseTTAFoldNA reporting encouraging accuracy for protein–NA interfaces and reasonably accurate RNA folds in many cases (8), and AlphaFold 3 extending structure prediction to biomolecular interactions including nucleic acids (9). Overall, tertiary-structure prediction is advancing rapidly, but robust generalization—especially to novel folds, long RNAs, and diverse experimental conditions—remains an open challenge highlighted across successive RNA-Puzzles rounds (10).

In practice, secondary structure is a particularly valuable intermediate representation: it is often more conserved than sequence, explains a large fraction of folding free energy, and serves as a routine basis for interpreting experiments and guiding downstream 3D modeling (11).

Because secondary structure can be represented as an *L* × *L* binary connectivity matrix, it naturally lends itself to both physics-based optimization and modern machine learning. Classical approaches are grounded in the thermodynamic nearest-neighbor model and dynamic programming, where structures are predicted by (approximate) free-energy minimization and related objectives; these ideas underlie widely used toolchains such as RNAstructure and ViennaRNA (12– 15). More recently, hybrid formulations have emerged that regularize or constrain learning using thermodynamic structure to improve robustness (16). In parallel, purely learning-based models have treated contact-map prediction as an “image-like” problem, achieving strong in-distribution accuracy with convolutional architectures (17, 18). Transformer-style and foundation-model approaches learn contextual representations from large unlabeled corpora and transfer them to structure prediction tasks (4). Finally, generative approaches have begun to model the distribution over plausible structures; diffusion-style formulations, for example, iteratively denoise a corrupted contact map to generate secondary structures while conditioning on sequence and learned embeddings (19).

Despite this progress, accurate RNA secondary-structure prediction remains difficult in realistic settings because the available sequence–structure datasets are frequently noisy, family-imbalanced, and non-stationary. Large-scale resources (e.g., bpRNA) are invaluable but can include labeling artifacts and uneven coverage across functional families (18). As a consequence, one-shot training often yields a trade-off: models can fit dominant families well yet generalize poorly to rare or functionally distinct RNAs, and performance degrades when the data regime shifts. When new families (or clusters) are introduced over time, naively continuing training can cause catastrophic forgetting, a well-known failure mode in sequential learning (20). These issues motivate moving beyond “train once on everything” toward methods that explicitly support continual integration of new RNA regimes without sacrificing previously learned competence.

To tackle these challenges, we propose RNAFoLBO (RNA Folding with Lifelong Bayesian Optimization), a continual-learning architecture for RNA secondary-structure prediction. RNAFOLBO treats latent RNA clusters (defined by clustering sequences in an embedding space) as a sequence of tasks and uses Lifelong Bayesian Optimization to orchestrate both model selection and hyperparameter tuning across heterogeneous predictors (UFold, RNA-FM, RNADiffFold), while preserving prior solutions rather than overwriting them. This yields a scalable alternative to repeated full retraining: as new clusters arrive, RNAFoLBO can allocate an appropriate model configuration for the new task while main-taining prior task-specific optima, thereby mitigating forgetting and improving stability. Empirically, across 15 latent clusters derived from RNAStrAlign-style benchmarks(15), RNAFOLBO achieves higher mean F1 and a substantially reduced performance range relative to strong single-model baselines, demonstrating that continual learning can provide both accuracy and robustness for practical, evolving RNA secondary-structure prediction.

A closely related methodological foundation for our pipeline is Lifelong Bayesian Optimization (LBO), an online multitask Bayesian optimization framework designed for settings where datasets (tasks) arrive sequentially and evolve over time (21). LBO casts model selection (and more generally, pipeline configuration) as black-box optimization and improves sample-efficiency by transferring information across tasks: instead of restarting Bayesian optimization from scratch for each new dataset, it reuses components of previously learned objective functions to warm-start and accelerate optimization on newly arriving tasks. This “lifelong” transfer is particularly appropriate for RNA secondary-structure prediction under distribution shift (e.g., new RNA families/clusters), where we want to rapidly adapt model/hyperparameter choices to a new cluster while leveraging experience from earlier clusters. We adopt this principle in RNAFoLBO by treating each latent RNA cluster as a task and using LBO-style transfer to guide sequential model selection and hyperparameter tuning.

## Methods

RNA secondary-structure prediction remains challenging despite decades of progress in thermodynamics-based algorithms and the rise of deep-learning architectures, and, more recently, generative approaches. Given its binary contact-matrix representation, deep-learning approaches typically treat secondary structure as a contact map prediction problem, using convolutional architectures that map sequence (and the derived pairwise features) to an *L* × *L* base-pair matrix. UFold, RNA-FM, and RNADiffFold, taken together, span three complementary and currently top-performing paradigms for RNA secondary-structure prediction, and therefore provide a fair and informative basis for evaluating our strategy. UFold (17) is a strong representative of supervised discriminative contact-map prediction: it directly maps sequence-derived features to an *L* × *L* base-pair matrix with an architecture optimized for accuracy and speed, and is widely used as a high-performing deep baseline for secondary structure inference. RNA-FM represents the rapidly growing class of foundation-model approaches for RNA: it learns general-purpose sequence representations from large unlabeled corpora and transfers them to downstream structure tasks, which is particularly relevant when data regimes are heterogeneous or when some clusters are label-poor—conditions that naturally arise in continual learning (4). RNADiffFold captures a third, increasingly influential direction: generative structured prediction via discrete diffusion, which explicitly models multi-modality by iteratively denoising a corrupted structure representation to produce plausible contact maps (19). By benchmarking against these three methods, we compare RNAFoLBO against (i) a best-in-class supervised predictor, (ii) a transfer-learning/foundation-model baseline, and (iii) a modern generative diffusion baseline—thereby covering the dominant modeling philosophies in the current literature while avoiding overfitting the evaluation to a single architectural family.

In fact, these architectures provide complementary inductive biases—convolutional “image-like” reasoning over contact maps (UFold), transformer-based contextual representations (RNA-FM), and generative diffusion modeling of multi-modal pairing (RNADiffFold).

In the following, we detail the proposed RNAFoLBO approach. In particular, section A formalizes the continual-learning problem setting for RNA secondary-structure prediction and defines how we construct the stream of tasks via latent RNA clustering. Section **??** describes the candidate prediction architectures considered in this work (UFold, RNA-FM, and RNADiffFold) and the unified training/evaluation interface used to compare them consistently across clusters. Section C then presents RNAFoLBO, our adaptation of the Lifelong Bayesian Optimization procedure that performs sequential model selection and hyperparameter tuning as new clusters arrive, enabling adaptation to distribution shift while mitigating catastrophic forgetting.

### A. RNAFOLBO algorithm overview

RNAFOLBO is a continual-learning pipeline that combines (i) latent-space clustering to define a meaningful stream of tasks from an imbalanced RNA dataset, and (ii) Lifelong Bayesian Optimization (LBO) to select, for each task/cluster, the best-performing folding architecture and hyperparameters, while transferring knowledge across clusters without overwriting prior solutions. Fig. 1 summarizes the end-to-end workflow and the information flow between components.

**Fig. 1.**
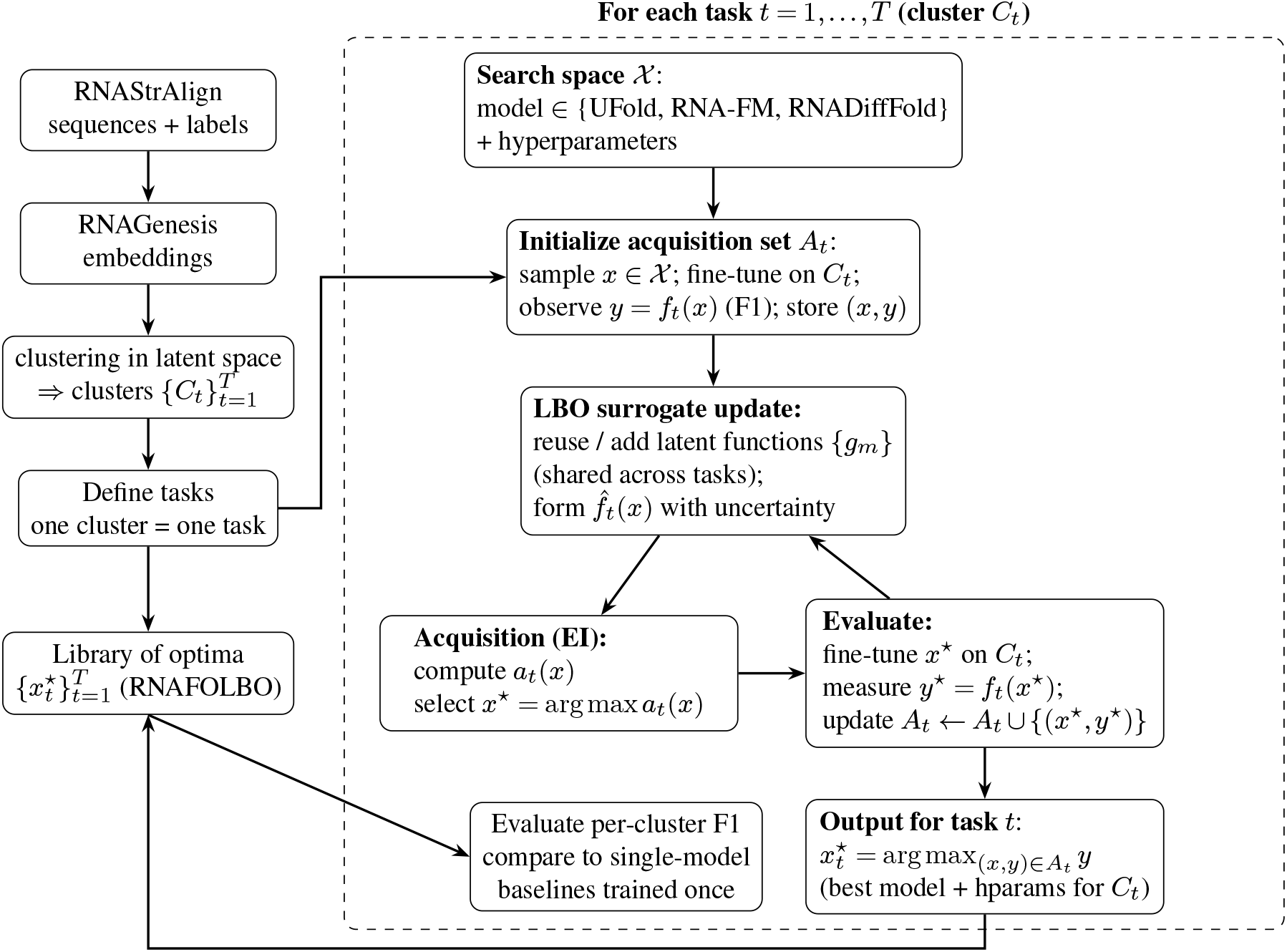
RNAFoLBO overview. Each latent RNA cluster (Cluster 0,…, Cluster 14) is treated as a task *t* with data 𝒟_*t*_. For that task, LBO defines a black-box objective *f*_*t*_(*x*), where *f*_*t*_(*x*) is the F1-score obtained after fine-tuning a candidate configuration *x ∈ X* (i.e., a specific model and hyperparameters, drawn from UFold, RNA-FM, RNADiffFold, …) on that cluster. Our Bayesian Optimization of *f*_*t*_ proceeds in three stages: *Initialization* (evaluate some initial configurations *x*), *Approximation* (learn a surrogate model of *f*_*t*_), and arg max_*x*_ *f*_*t*_(*x*) (select the best-performing configuration for that cluster). The surrogate for *f*_*t*_ is constructed using a set of latent basis functions *g*_*m*_, which are shared across clusters and allocated via an Indian Buffet Process. These shared functions enable knowledge transfer across biologically similar clusters instead of relearning from scratch. Running this procedure for each cluster *t* yields an optimal configuration 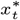 (shown as *x*_0_, *x*_1_, *x*_2_, …, *x*_14_): one specialized model and associated hyperparameters per cluster, rather than a single global model. This avoids catastrophic forgetting and enables scalable, cluster-specific RNA secondary-structure prediction.

Task construction via embeddings and clustering. The first component transforms each RNA sequence into a contextual embedding using RNAGenesis, which couples a Transformer encoder with a Q-former to extract rich sequence features and induces a structured latent space through a diffusion process. We then apply clustering in this latent space to obtain groups 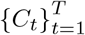 (classes) that are substantially more balanced than the original functional-family partition, and we treat each cluster *C*_*t*_ as a task in the continual-learning sense (Fig. 1, left branch). This step is essential because LBO requires a stream of tasks with enough samples per task to support within-task fine-tuning and evaluation; clustering in the RNAGenesis latent space provides exactly this task definition and enables scalable continual updates as new RNA regimes appear.

Continual learning by LBO over heterogeneous folding models. Once tasks are defined, RNAFOLBO runs an LBO loop independently for each cluster *C*_*t*_ to optimize a black-box objective *f*_*t*_ (*x*) : *X* → ℝ, where *x* ∈ *X* denotes a configuration (model choice and associated hyperparameters) drawn from a heterogeneous candidate set (e.g., UFold, RNA-FM, RNAD-iffFold), and *f*_*t*_ (*x*) is the F1 score obtained after fine-tuning that configuration on the data in cluster *C*_*t*_ (Fig. 1, middle branch). Algorithm 1 provides the corresponding pseudocode: we first initialize an acquisition set *A*_*t*_ by evaluating a small number of configurations *x* (fine-tune on *C*_*t*_, measure F1, store (*x, f*_*t*_ (*x*)), then repeat for *N*_acq_ acquisition rounds: update a task-specific surrogate 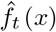, select the next configuration *x*^*^ by maximizing an acquisition function *a*_*t*_ (*x*) (Expected Improvement, EI), evaluate *x*^*^ on *C*_*t*_, and aug-ment *A*_*t*_. The output of task *t* is the best observed configu-ration 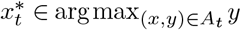 (Algorithm 1; Fig. 1 “Output for task *t*”).

#### Algorithm 1

Lifelong Bayesian Optimization for RNA secondary-structure prediction

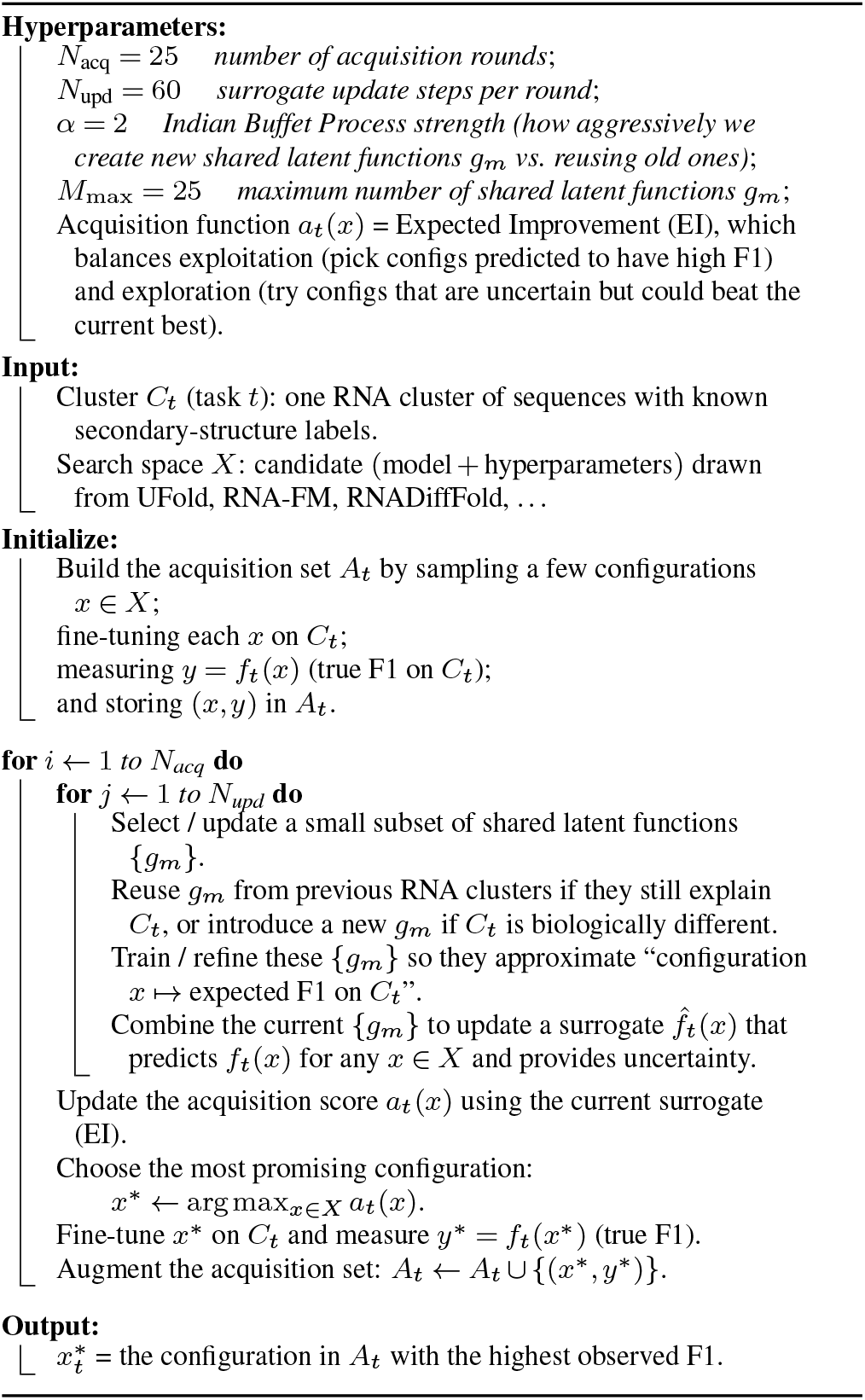

### B. Determining the Tasks

In continual learning, the notion of a “task” is only useful if each task provides enough samples to support learning and evaluation; otherwise the problem collapses into a few-shot regime that is ill-suited to RNA secondary-structure prediction. We therefore redefine tasks globally by constructing a stream of size-balanced, latent RNA clusters. In fact, in several data sets for RNA, defining tasks directly from functional families would lead to a strongly imbalanced learning problem. Some tasks would contain very few samples, effectively turning the problem into a *few-shot learning* scenario. However, few-shot learning is generally poorly suited for RNA secondary-structure prediction, as these models typically require a sufficient number of labeled examples to reliably learn sequence–structure relationships and base-pairing patterns. To avoid this issue and obtain tasks of comparable size, we instead redefine tasks by clustering sequences in a latent representation space.

Let *x*_*i*_ denote an RNA sequence. We compute a contextual representation *ϕ* (*x*_*i*_) ∈ ℝ^*d*^ using RNAGenesis (22), a model specialized for contextual representation learning and latent-space structuring. RNAGenesis couples a Transformer encoder with a Q-former module to extract high-resolution contextual features from each sequence; the resulting embedding is then processed through a diffusion-based latent construction mechanism, where the diffusion input defines the latent space used for downstream operations (including generation of plausible sequences). In our pipeline, these embeddings serve as a geometry-preserving representation in which similarity reflects sequence context beyond simple edit distance, enabling task definition that is not tied to the original, imbalanced family labels.

Given embeddings 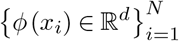, we evaluated two complementary clustering methods to define tasks, *k*-means and spectral clustering. *k*-means is computationally efficient and provides a direct knob on the granularity of each task (controlling the target number of clusters *T*). Such bound is useful in ensuring each task has sufficient samples. We considered spectral clustering to capture potentially non-convex cluster structure This approach is more expressive than *k*-means in the original space because the Laplacian eigenvectors can separate clusters connected by manifold-like geometry and can therefore better respect the intrinsic topology of *ϕ* (·). In both cases, each cluster *C*_*t*_ defines one task for the continual-learning stage.

The clustering step is not only a preprocessing choice; it makes the downstream optimization problem well-posed. For each task/cluster *C*_*t*_, we define a task-specific dataset 𝒟_*t*_ on which candidate secondary-structure predictors can be fine-tuned and evaluated, producing a task objective *f*_*t*_ (*x*) (F1 score) for any configuration *x* (model choice + hyperparameters). As shown in the workflow figure (Fig. 1), the embeddings-and-clustering block outputs the stream of tasks {*C*_*t*_}, which drives the per-task LBO loop (initialization, surrogate update, acquisition, evaluation, and task optimum). Running this process across tasks yields a library of cluster-specific optima rather than a single global model choice dominated by the most frequent regimes.

### C. Continual Learning with Alternative Folders

As previously mentioned, Lifelong Bayesian Optimization (LBO (21)) allows RNAFoLBO to leverage multiple folding algorithms while at the same time addressing the issues of scalability. Specifically, for each task *t*, and corresponding cluster *C*_*t*_ identified during the clustering step (section B), we can observe the F1 score associated with a specific model *m* ∈ ℳ and hyperparameter choice *θ*_*m*_ ∈ Θ_*m*_ (generally each model may have a different hyperparameter space), by training the model for the corresponding data set 𝒟_*t*_. Arguably, such training is very expensive, so, under the LBO architecture, we learn a surrogate as a cheaper function to predict the F1 score under configurations not tested. Specifically, the learned surrogate maps a model, hyperparameter selection to the F1 score, namely *f*_*t*_ : ℳ Θ ↦ ℝ, where Θ = {Θ_*m*_}. For convenience, we will refer to the product space of model and hyperparameters as the search space 𝒳. Noticeably, this is a mixed categorical (*model* choice) and continuous *hyperparameters* search space. The Bayesian optimization search is then called for each task *t* to sequentially generate candidate locations *x* ∈ 𝒳 is a way that improves the true F1 score. So, different from continual learning, we can change the model to employ at each task not just adapt the parametrization.

As a result of the LBO step we obtain as many combinations of models and associated hyperparameters as many tasks we have. This is an important aspect of LBO: since each task retains its own best configuration, catastrophic forgetting is drastically reduced. In fact, adding a new task does not over-write the model used by prior tasks. In the LBO formulation, *f*_*t*_ is further expressed through a sparse combination of shared latent functions *g*_*m*_ (small neural networks) that are learned across tasks; this allows knowledge transfer when a new task resembles a previous one while remaining flexible when it differs (see Fig. 1 and Alg. 1).

## Results

### Sample Data

The dataset selected for this study is the RNAStrAlign (Fig. 2), a widely used benchmark for RNA secondary-structure prediction, which provides both the sequence and the annotated secondary structure of RNA molecules. The sequences are organized into *functional RNA families*, grouping molecules that share similar biological roles and structural characteristics, which facilitates comparative analyses across related families. In fact, Fig. 2 shows how imbalanced is the distribution of sequences across *functional families*.

**Fig. 2.**
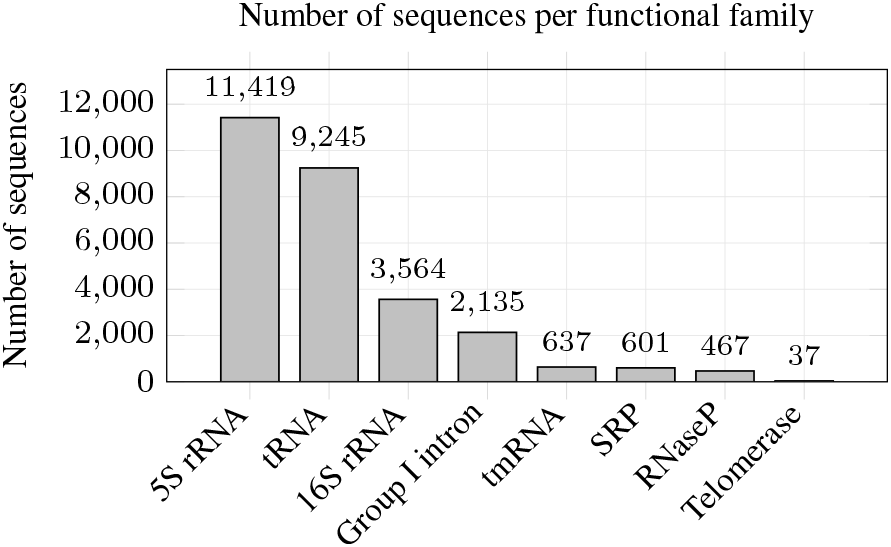
Distribution of RNA sequences across functional families within the RNAS-trAlign dataset.

While families such as 5S rRNA and tRNA contain thousands of sequences, others such as Telomerase contain only a few dozen. This strong imbalance motivates redefining tasks through latent-space clustering rather than directly using functional families.

### Benchmarks

We used RNAGenesis (22), a model specialized in contextual representation learning, to cluster our dataset. In this experimental analysis, for each cluster, we test three folding approaches, i.e., | ℳ |= 3, UFold, RNA-FM, and RNADiffFold, which we chose based on the reported performance. Concerning the set Θ = {Θ_*m*_}, we decided to fix the set cardinality across the different models (not the same hyperparameters). Specifically, while the learning rate *l*_*m*_ has the same semantics across the three models, the second hyperparameter *λ*_*m*_ plays a regularizing role, discouraging memorization of a single cluster and promoting generalization to sequences within or near that cluster. Below, we list the hyperparameters with the associated support used to execute LBO over incoming tasks (resulting from the embedding-driven clustering in Section B):

- *m* = 1, UFold: *l*_1_ ∈ [10^−5^, 3 · 10^−4^], we used the post-processing strength as regularizing hyperparameter *λ*_1_ *∈* [1, 5] (post-processing strength);
- *m* = 2, RNA-FM: *l*_2_ ∈ [3 · 10^−6^, 3 · 10^−4^], we used the scheduler warm up ratio as regularizing hyperparameter *λ*_2_ *∈* [0.03, 0.12];
- *m* = 3, RNADiffFold: *l*_3_ ∈[5·10^−6^, 3·10^−4^], we chose the loss weight as regularizer *λ*_3_ *∈* [0.2, 2.0]

To establish a comparison with standard deep learning, we also train each model “conventionally”, i.e., against the full dataset, then evaluate these “globally trained” models on each of the *T* tasks produced by the clustering step based on the RNAGenesis embedding. For training, we chose to use 80% of the data for training and 20% for testing. Indeed, we do not include a separate validation step, as its primary purpose is to optimize hyperparameters, which is already handled by the LBO strategy.

### Performance Metrics

Our analysis focuses primarily on the *F* 1 score. Specifically, we compare the F1-score of RNAFoLBO (averaged across the *T* clusters) with that of each baseline model “trained globally”. The idea is to show the overall effectiveness of the CL-based approach compared to traditional training approaches. We also analyze the distribution of the RNAFoLBO F1-score across tasks, comparing for each cluster the performance of our approach with that of the global models evaluated on that same cluster. This finer-grained analysis identifies scenarios where specialized task-based learning provides a significant advantage.

**In fact, the F1-score of a model trained and evaluated on the entire dataset should differ from the mean of the cluster-wise F1-scores, since the F1-score is not a linear metric. However, because our clusters exhibit the same order of magnitude, these two quantities are nearly equal, enabling a direct comparison between macro- and micro-level approaches**.

We report the performance range (the difference between the maximum and minimum F1-score achieved across tasks) for each method. This metric serves as a simple but informative indicator of model stability, where a smaller range indicates more consistent performance across different RNA regimes.

### D. Understanding the effect of the RNAGenesis latent space on families re-clustering

In order to better understand the effect of the embedding and the quality of the clusters homogeneity, we performed a qualitative analysis of the latent structure using several dimensionality-reduction techniques. We explored several dimensionality-reduction techniques, including PCA, t-SNE, and UMAP. Under each projection method, hyperparameters were first explored in order to obtain visually informative embeddings. The clustered sequences were then projected into the reduced space and colored according to their cluster assignment. These visualizations provide qualitative evidence that the structure induced by the latent space does increase homogeneity in the nanoparticles and that the obtained clusters are reasonably well separated.

We eventually focused on UMAP (Uniform Manifold Approximation and Projection) projections (23). This technique is well-suited for preserving both local and global structure in high-dimensional embeddings, making it ideal for assessing whether our latent clusters correspond to meaningful biological groupings. We systematically varied UMAP’s key hyperparameters, the minimum distance and the number of neighbors to identify the most informative projections: (i) the minimum distance (`min_dist`) controls the density of the projected points. Smaller values produce more compact clusters, while larger values preserve more global structure at the cost of local separation; (ii) the number of neighbors (`n_neighbors`) determines the size of the local neighborhood used to construct the manifold. Lower values emphasize fine-grained local structure, whereas higher values capture broader topological patterns.

Based on the visual inspection of the projections, we selected `min_dist` = 1.0 and `n_neighbors` = 2 for our experiments. This configuration produces a spread and relatively uniform embedding, avoiding the introduction of artificial cluster structure while preserving local neighborhood relationships. Such a representation is particularly suitable for visualizing the cluster assignments obtained from applying *k*-means in the latent space, as it allows the evolution and spatial organization of these clusters to be observed without imposing additional separation in the projection (see Fig. 4). Thus, Fig. 5 displays how our clustering looks like when sequences are projected using UMAP. We can see that, visually, even if the distribution of clusters appears diffuse, all clusters are noticeable. Fig. 6 shows how the sequences initially assigned to a single family end up being split across the cluster, with the families associated with highest number of sequences being split across several of the embedding-induced clusters, This has an important balancing effect: in Fig. 2 the first two families contain ∼ 75.5% of the data, while the 2 most populated clusters we obtain contain 40% of the data.

**Fig. 3.**
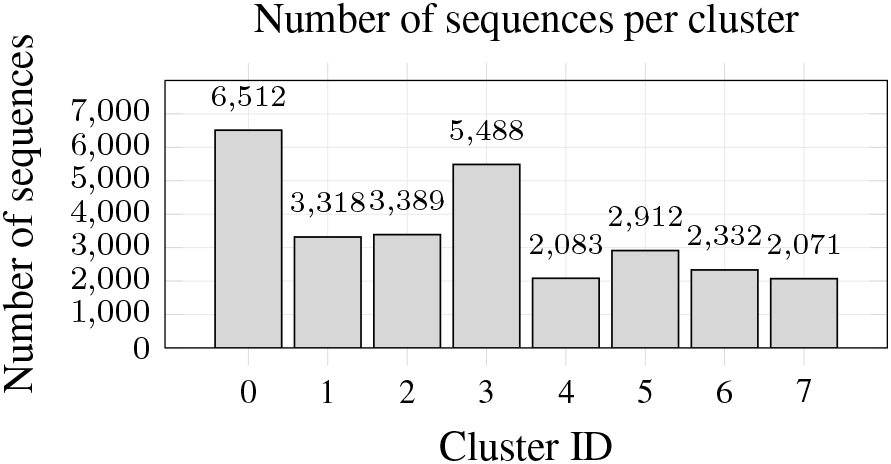
Latent-space clustering balances sample sizes. Functional families in the original dataset are highly imbalanced, whereas clustering sequences in the latent space learned by RNAGenesis—a model specialized in contextual representation learning and latent-space structuring— using K-means yields clusters with the same order of magnitude, enabling fairer training and evaluation.

**Fig. 4.**
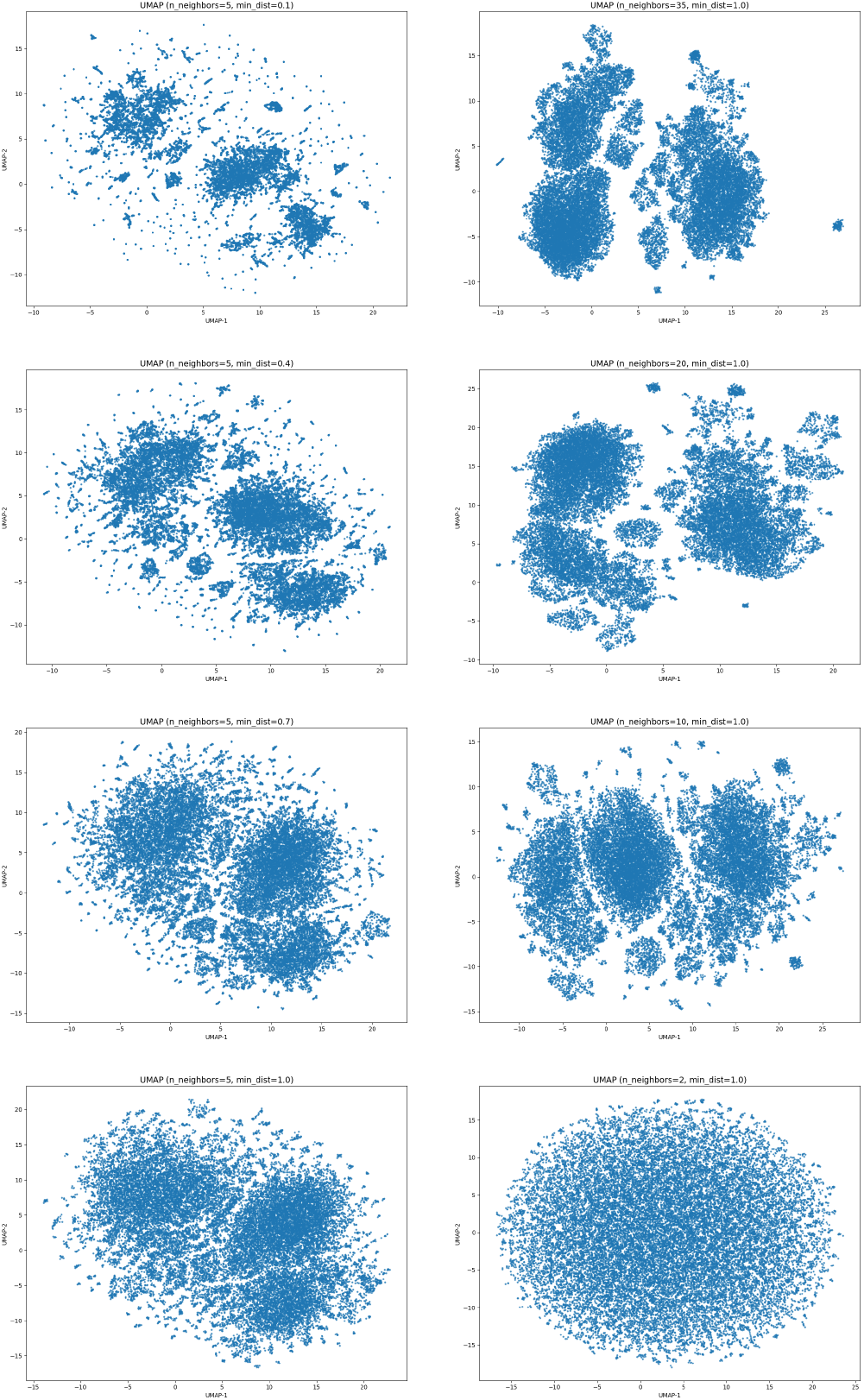
Effect of UMAP hyperparameters. The left column varies the `min_dist` parameter (0.1–1.0) while keeping `n_neighbors` fixed, illustrating how increasing `min_dist` progressively relaxes cluster compactness and emphasizes global structure. The right column varies `n_neighbors` (35–2), showing the transition from broader manifold connectivity to highly local neighborhood structure as the neighborhood size decreases. Together, these projections illustrate how UMAP balances local cluster separation and global manifold preservation.

**Fig. 5.**
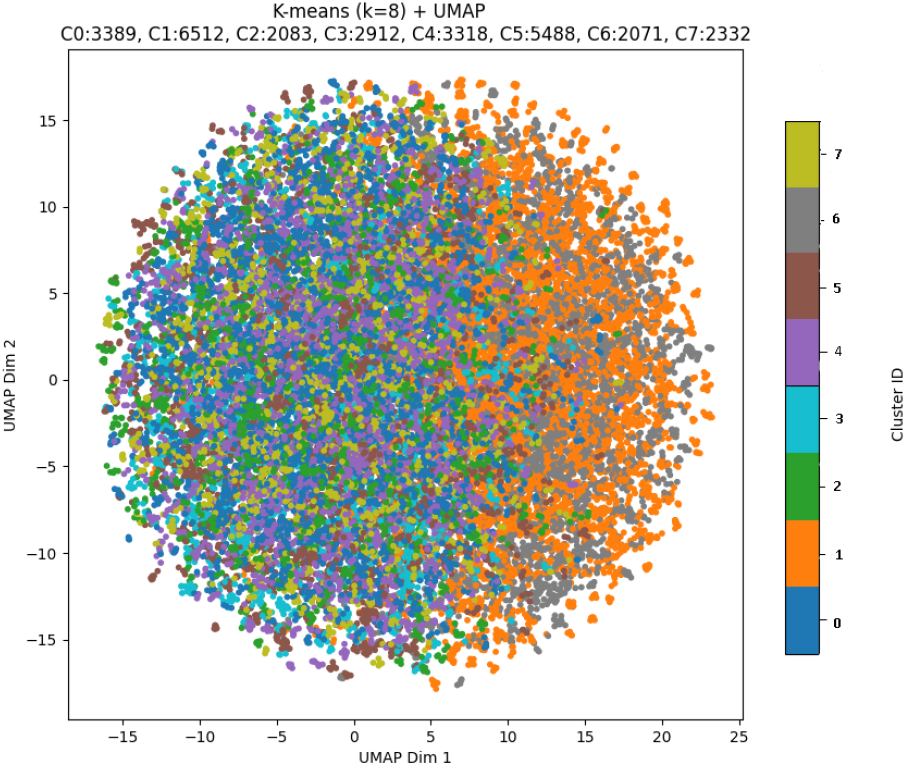
K-means clustering in latent space. Sequences are projected using UMAP (`min_dist = 0.1, n_neighbors` = 2) and colored according to the 8 clusters obtained with K-means in the RNAGenesis latent space. The resulting clusters are relatively balanced and all clusters are noticeable.

**Fig. 6.**
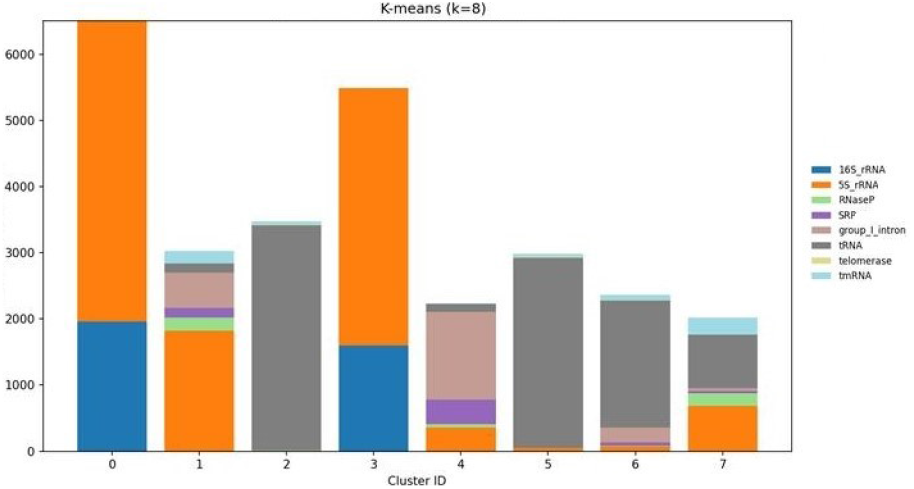
Famillies repartition across our clustering. This figure display the repartition of the functional families across our clustering. In particular, we can notice that the heaviest families (5s rRNA and tRNA) are distributed in several of our clusters, thus balancing the distribution of sequences

#### Choice of the number of clusters

As mentioned in Section B we tested both *k*-means and spectral clustering. Specifically, for each clustering method, we tested for *T* = 1,…, 20, and selected the number of clusters guaranteeing the resulting partition to remain reasonably balanced. After comparing the different clustering strategies, *k*-means was found to produce the most balanced and interpretable clusters and was therefore selected for the remainder of the experiments.

The next step was to determine the number of clusters. This choice involves a trade-off: a sufficiently large number of clusters allows the continual-learning framework to exploit transfer across tasks, while too many clusters may lead to clusters that are too small for reliable training.

In our experiments, a total of *T* = 15 clusters provided a good balance between these two constraints.

### E. Performance comparison

The different F1-scores discussed above are reported in Table 1 and are also presented graphically in Figure 7.

**Table 1.** Per-cluster F1-scores for the 15 latent clusters (0–14). Each row is one cluster. RNAFoLBO reports, for each cluster, the best combination of model and hyperparameters chosen by RNAFoLBO ; UFold, RNA-FM, and RNADiffFold are single-model baselines trained once on the full dataset. In the RNAFoLBO column, blue values are clusters where RNADiffFold was selected (all clusters except 4,12 and 13), and green values are clusters where RNA-FM was selected (clusters 4, 12, and 13). The last two rows summarize the mean F1 and the range (max–min) per method.

| $t$ | FoLBO | UFold | FM(★) | DiffFold(★★) |
| --- | --- | --- | --- | --- |
| 0 | 0.911(★★) | 0.861 | 0.746 | 0.903 |
| 1 | 0.965(★★) | 0.570 | 0.809 | 0.858 |
| 2 | 0.931(★★) | 0.604 | 0.759 | 0.934 |
| 3 | 0.981(★★) | 0.898 | 0.954 | 0.958 |
| 4 | 0.821(★) | 0.506 | 0.768 | 0.577 |
| 5 | 0.886(★★) | 0.562 | 0.777 | 0.880 |
| 6 | 0.937(★★) | 0.615 | 0.689 | 0.940 |
| 7 | 0.987(★★) | 0.512 | 0.940 | 0.960 |
| 8 | 0.986(★★) | 0.586 | 0.987 | 0.963 |
| 9 | 0.942(★★) | 0.609 | 0.741 | 0.944 |
| 10 | 0.901(★★) | 0.585 | 0.779 | 0.901 |
| 11 | 0.981(★★) | 0.554 | 0.984 | 0.950 |
| 12 | 0.810(★) | 0.390 | 0.769 | 0.458 |
| 13 | 0.983(★) | 0.595 | 0.985 | 0.963 |
| 14 | 0.946(★★) | 0.597 | 0.774 | 0.944 |
| $\mu$ | 0.931 | 0.602 | 0.830 | 0.875 |
| $\vert \Delta F1\vert $ | 0.177 | 0.508 | 0.298 | 0.505 |

**Fig. 7.**
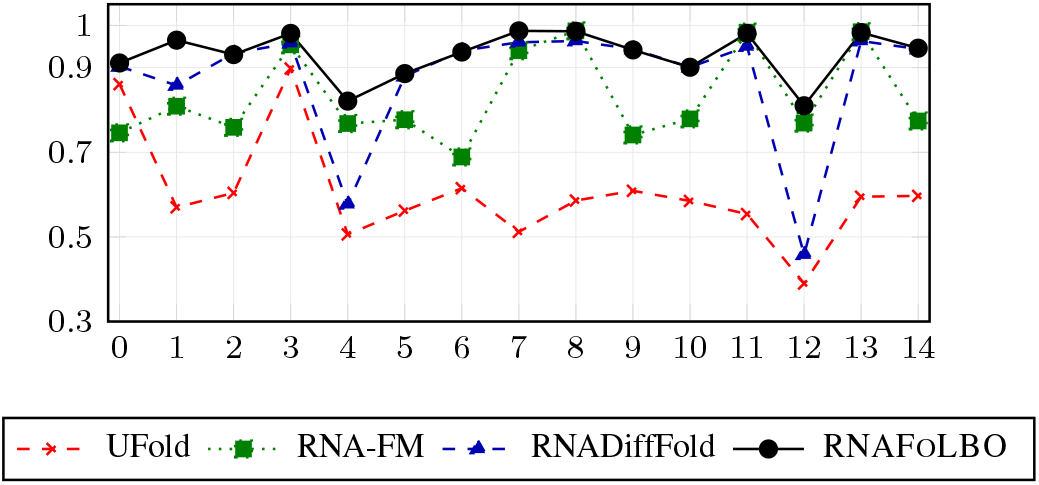
Per-cluster F1-scores (clusters 0–14). The proposed RNAFoLBO is the best-per-cluster configuration chosen by Lifelong Bayesian Optimization. UFold, RNA-FM, and RNADiffFold are single-model baselines trained once on the full dataset, then evaluated on each cluster.

As we can see, the RNAFoLBO F1 curve largely upper-bounds those of the individual deep-learning baselines. Looking at per-cluster values, RNADiffFold emerges as the most accurate model (selected 12*/*15 times by RNAFoLBO), yet it is not the most stable (larger range). RNAFoLBO thus exploits RNADiffFold’s strengths—consistent with external benchmarks—while addressing its weaknesses (clusters 4, 12, and 13) by switching in those cases to a slightly less accurate but more stable model specialized in contextual representation learning: RNA-FM. This strategy allows RNAFoLBO to achieve a higher average performance than the stand-alone baselines F1 = 0.931 and greater stability, with a range of 0.177 demonstrating its robustness.

### F. RNAFOLBO sequence-by-sequence comparison

To complement the aggregate and per-cluster *F* 1 analysis, we provide a sequence-level qualitative comparison that makes the nature of the errors (and corrections) visually explicit. Fig. 8-9 report, for two representative sequences each, (i) a 2D secondary-structure drawing and (ii) the associated connectivity-map view. Each figure is organized by columns: Ground Truth, RNA-FM (classic one-shot training), RNADiffFold (classic one-shot training), and RNAFoLBO (continual-learning pipeline). The connectivity-map panels use a common legend: the ground-truth base pairs are shown in blue, while predictions are decomposed into true positives (green), false positives (red), and false negatives (orange), allowing direct visual diagnosis of which helices are recovered, spurious, or missed.

**Fig. 8.**
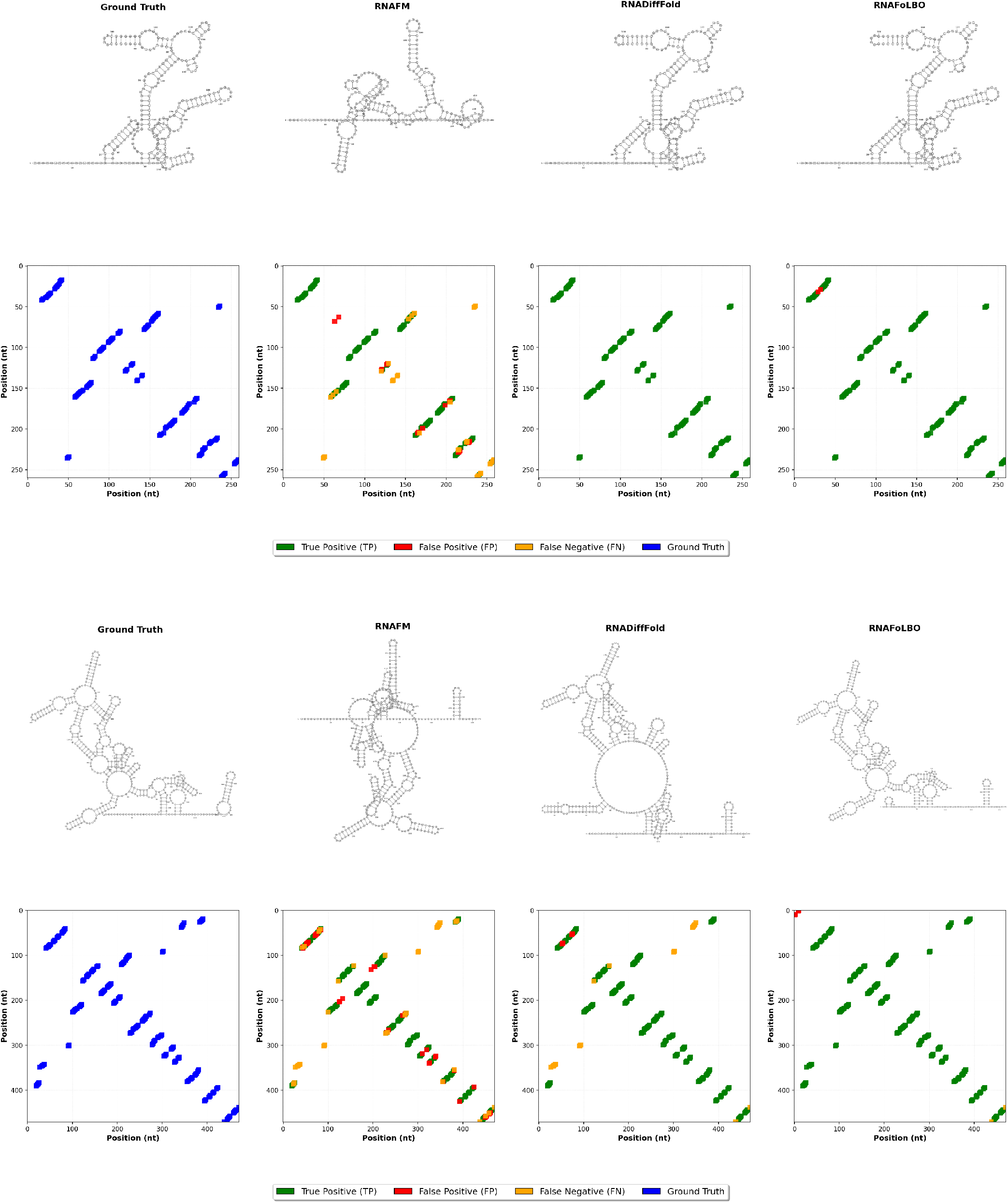
Those sequences (X63823 and AM051262) are from cluster where the retained model is RNADiffFold. For each sequence, you can find a visual representation and the contact map of the ground truth as well as the predictions of RNADiffFold and RNAFM (classic learning), with also RNAFoLBO (Continual learning)

Fig. 8 reports results for two sequences (X63823 and AM051262) sampled from a cluster where the cluster-retained model selected by RNAFoLBO is RNADiffFold. Importantly, these examples illustrate that RNAFoLBO is not merely “reprinting” the vanilla RNADiffFold prediction: even when RNADiffFold is the retained architecture for the cluster, RNAFoLBO ‘s cluster-specific configuration (i.e., the LBO-selected hyperparameters and the cluster-specialized fine-tuning) can materially change the predicted structure. This is clearest in the second row of Fig. 8, where both one-shot baselines exhibit noticeable failure modes in the global topology (RNA-FM produces a heavily distorted fold and RNADiffFold introduces a large erroneous loop/bulge relative to the ground truth), whereas RNAFoLBO recovers a structure that is visually closest to the reference, with a cleaner alignment of the dominant helical segments and fewer gross topological artifacts. In connectivity-map terms, the RNAFoLBO column shows a denser concentration of true positives along the main diagonals corresponding to the ground-truth stems, while suppressing the spurious pairings that drive the baseline failures.

Similarly, Fig. 9 repeats the same diagnostic layout for two sequences (P00797 and Gal.c.trnL) drawn from a cluster where the retained model is RNA-FM. Again, the key point is that RNAFoLBO ‘s improvement is not explained by simply returning the “best baseline” unchanged. In the second row of Fig. 9, both classic baselines fail in complementary ways (RNA-FM misses substantial portions of the reference pairing pattern, while RNADiffFold introduces systematic off-target pairings), producing contact maps with substantial orange (misses) and red (spurious) regions. By contrast, RNAFoLBO produces a prediction that is closer to the ground truth drawing and exhibits a more faithful recovery of the dominant stems in the connectivity—i.e., more green aligned with the blue reference and fewer large-scale structural distortions.

**Fig. 9.**
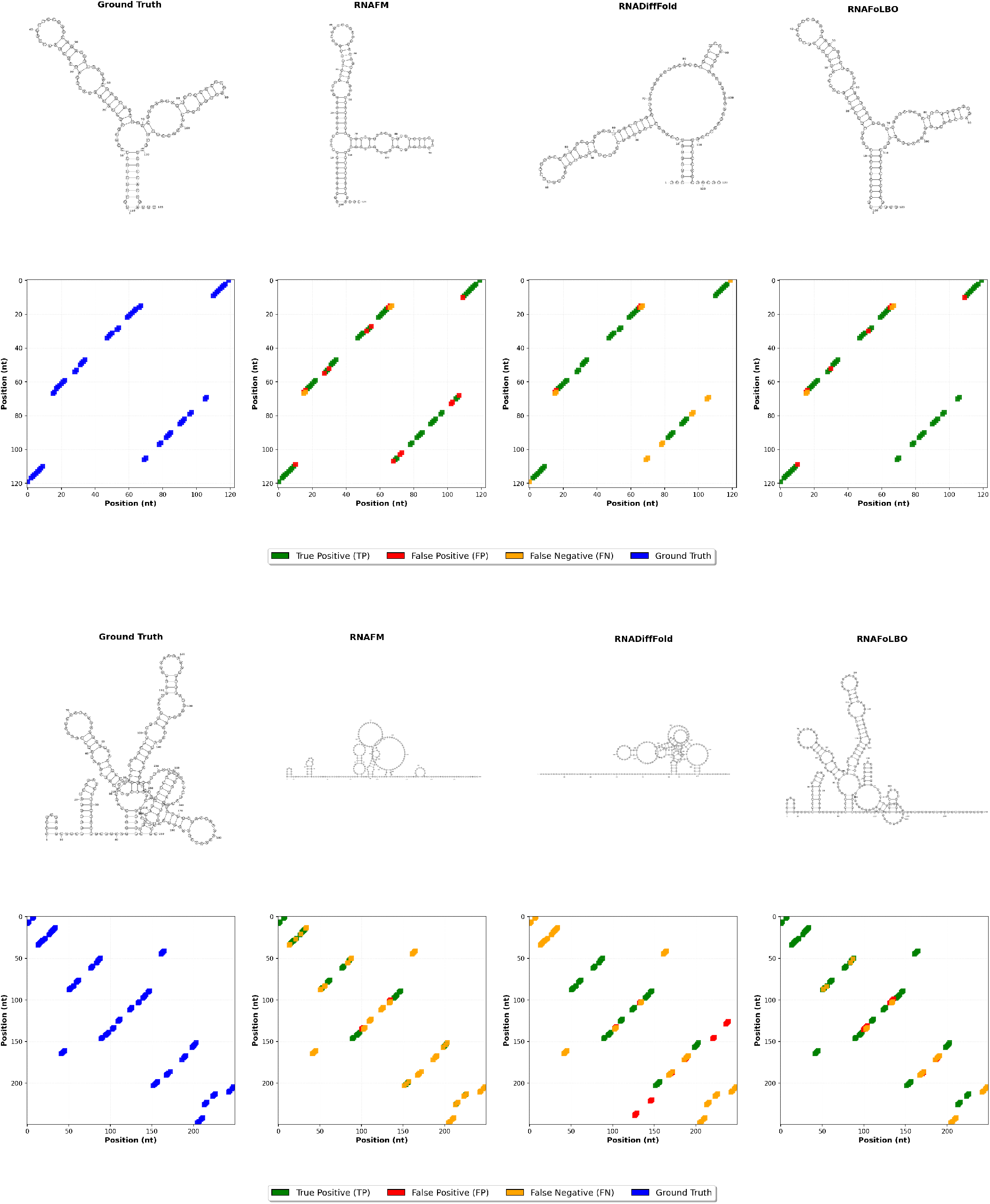
Those sequences (P00797 and Gal.c.trnL) are from cluster where the retained model is RNAFM. For each sequence, you can find a visual representation and the contact map of the ground truth as well as the predictions of RNADiffFold and RNAFM (classic learning), with also RNAFoLBO (Continual learning)

Overall, Figs. 8-9 provide qualitative evidence for two distinct mechanisms behind RNAFOLBO’s gains: (i) cluster-appropriate model selection (RNA-FM vs. RNADiffFold depending on the cluster), and (ii) within-architecture improvements driven by LBO’s cluster-specific configuration search—so that even in clusters where the retained architecture matches a strong baseline, RNAFOLBO can still correct failure cases and produce structures that are visibly closer to the ground truth rather than merely reproducing the baseline output.

## Discussion

### Possible extensions of the proposed RNAFoLBO approach

While the empirical results show the performance of RNAFoLBO against the families within the data set of choice, two practical scenarios follow: (i) new incoming RNA sequence, (ii) new incoming family. If we need to predict the structure of a single RNA sequence, we first assign the sequence to the closest latent cluster (e.g., by distance in the RNAGenesis embedding space). We then use the RNAFoLBO -selected model for that cluster to produce the most reliable secondary-structure prediction. If we want to add a new family of sequences, it becomes a new task and does not affect previously defined tasks. In RNAFoLBO, each task (*t*) keeps its own optimal configuration 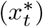 and trained weights. The black-box objective is modeled as a sparse combination of shared latent functions ({*g*_*m*_}); when a new task arrives, the prior underling the LBO algorithm selects a small subset (*S*_*t*+1_) of these ({*g*_*m*_}) (and instantiates new ones if needed). This sharing provides positive transfer, while sparsity limits negative transfer. Crucially, since the ({*g*_*m*_}) are only used to guide acquisition (model/hyperparameter selection) and we do not overwrite the stored (*x*^*^) models, performance on earlier tasks remains unchanged (no catastrophic forgetting).

#### Computational Limitations

A key limitation of our approach is its computational overhead. To approximate each task-specific objective *f*_*t*_, RNAFoLBO must evaluate mul-tiple candidates *x* ∈ *X* ; each evaluation entails fine-tuning the chosen model on cluster *t* and measuring the associated F1 score. In our setup, the classical baselines required only 7 epochs of fine-tuning per model on the full dataset, whereas RNAFoLBO —because it repeatedly fine-tunes across many candidates and across the 15 clusters—accumulated 2625 epochs in total, distributed among UFold, RNA-FM and RNADiffFold. Although the surrogate updates (shared latent functions and acquisition) are lightweight compared with training, the wall-clock and GPU budget are substantially higher. As usual, there is a trade-off between *performance* (higher and more stable with LBO) and *cost* (greater compute and tuning time).

#### Domain compatibility (intersection of domains of definition)

To compare models fairly, we restrict ourselves to the overlap of inputs that all models can handle—the intersection of the model’s domains of definition. In practice, each model has its own operational limits (e.g., maximum sequence length, tokenization/alphabet, input format). We therefore keep only sequences that satisfy every model’s requirements at once. In our setup, the tightest shared constraint is length, so we cap sequences at 512 nucleotides and discard longer ones. RNAFoLBO does not have any limitation in this sense.

## Conclusions

We propose RNAFoLBO to address the limitations of classical deep-learning approaches to RNA secondary-structure prediction—namely their sensitivity to noisy, family-imbalanced datasets, reduced out-of-distribution generalization, and catastrophic forgetting when new data regimes are added—by adopting a continual learning method. Because continual learning relies on multiple tasks and functional families are highly imbalanced, we redefine tasks by clustering the entire dataset in the latent space of RNAGenesis, a model specialized in contextual representation learning and RNA sequence generation (i.e., latent-space structuring). We also filter sequences to the *overlap of inputs all models can process*, so that UFold, RNA-FM, and RNADiffFold can be evaluated on the exact same examples (in practice, this entails discarding sequences longer than 512 nucleotides).

With tasks defined, we adapt LBO to approximate, for each task *t*, the black-box objective *f*_*t*_ : *X* → ℝ, where *X* is the joint space of *model choice* and *hyperparameters*, and *f*_*t*_(*x*) returns the F1 score obtained after fine-tuning configuration *x* on task *t*. Bayesian optimization (via a shared surrogate and an acquisition function) explores *X*, and we retain per cluster the single best model together with its optimal hyperparameters (UFold, RNA-FM, or RNADiffFold).

RNAFoLBO delivers both higher average performance and greater stability than single-model deep-learning baselines, and it scales more gracefully: adding a new task does not degrade performance on previous tasks because each task preserves its own selected configuration, while the shared surrogate transfers knowledge through latent functions without overwriting earlier solutions—thereby mitigating catastrophic forgetting.

## Bibliography

1. Laura Teodori, Marjan Omer, and Jørgen Kjems. Rna nanostructures for targeted drug delivery and imaging. RNA biology, 21(1):391–409, 2024.

2. Justin T Low and Kevin M Weeks. Shape-directed rna secondary structure prediction. Methods, 52(2):150–158, 2010.

3. Peixuan Guo. The emerging field of rna nanotechnology. Nature nanotechnology, 5(12):833–842, 2010.

4. J. Chen et al. Interpretable rna foundation model from unannotated data (rna-fm). bioRxiv, 2022. doi: 10.1101/2022.08.06.503062.

5. Hannah K Wayment-Steele, Wipapat Kladwang, Alexandra I Strom, Jeehyung Lee, Adrien Treuille, Alex Becka, Eterna Participants, and Rhiju Das. Rna secondary structure packages evaluated and improved by high-throughput experiments. Nature methods, 19(10):1234– 1242, 2022.

6. José Almeida Cruz, Marc-Frédérick Blanchet, Michal Boniecki, Janusz M Bujnicki, Shi-Jie Chen, Song Cao, Rhiju Das, Feng Ding, Nikolay V Dokholyan, Samuel Coulbourn Flores, et al. Rna-puzzles: a casp-like evaluation of rna three-dimensional structure prediction. Rna, 18(4):610–625, 2012.

7. Andrew Martin Watkins, Ramya Rangan, and Rhiju Das. Farfar2: improved de novo rosetta prediction of complex global rna folds. Structure, 28(8):963–976, 2020.

8. Minkyung Baek, Ryan McHugh, Ivan Anishchenko, Hanlun Jiang, David Baker, and Frank DiMaio. Accurate prediction of protein–nucleic acid complexes using rosettafoldna. Nature methods, 21(1):117–121, 2024.

9. Josh Abramson, Jonas Adler, Jack Dunger, Richard Evans, Tim Green, Alexander Pritzel, Olaf Ronneberger, Lindsay Willmore, Andrew J Ballard, Joshua Bambrick, et al. Accurate structure prediction of biomolecular interactions with alphafold 3. Nature, 630(8016):493– 500, 2024.

10. Fan Bu, Yagoub Adam, Ryszard W Adamiak, Maciej Antczak, Belisa Rebeca H de Aquino, Nagendar Goud Badepally, Robert T Batey, Eugene F Baulin, Pawel Boinski, Michal J Boniecki, et al. Rna-puzzles round v: blind predictions of 23 rna structures. Nature methods, 22(2):399–411, 2025.

11. J. Fallmann, S. Will, J. Engelhardt, et al. Recent advances in rna folding. Journal of Biotechnology, 2017. doi: 10.1016/j.jbiotec.2017.07.007.

12. Michael Zuker and David Sankoff. Rna secondary structures and their prediction. Bulletin of Mathematical Biology, 1984. doi: 10.1007/BF02459506.

13. Jessica S. Reuter and David H. Mathews. Rnastructure: software for rna secondary structure prediction and analysis. BMC Bioinformatics, 2010. doi: 10.1186/1471-2105-11-129.

14. Ronny Lorenz, Stephan H. F. Bernhart, Christian Höner zu Siederdissen, et al. Viennarna package 2.0. Algorithms for Molecular Biology, 2011. doi: 10.1186/1748-7188-6-26.

15. Zhen Tan, Ying Fu, Gunjan Sharma, and David H. Mathews. Turbofold ii: Rna structural alignment and secondary structure prediction informed by multiple homologs. Nucleic Acids Research, 2017. doi: 10.1093/nar/gkx815.

16. Kengo Sato, Masanori Akiyama, and Yasubumi Sakakibara. Rna secondary structure prediction using deep learning with thermodynamic integration. Nature Communications, 2021. doi: 10.1038/s41467-021-21194-4.

17. L. Fu, Y. Cao, J. Wu, et al. Ufold: fast and accurate rna secondary structure prediction with deep learning. Nucleic Acids Research, 2022. doi: 10.1093/nar/gkab1074.

18. Parinaz Danaee, M. Rouches, M. Wiley, et al. bprna: large-scale automated annotation and analysis of rna secondary structure. Nucleic Acids Research, 2018. doi: 10.1093/nar/gky285.

19. Z. Wang, Y. Feng, Q. Tian, et al. Rnadifffold: generative rna secondary structure prediction using discrete diffusion models. Briefings in Bioinformatics, 2024. doi: 10.1093/bib/bbae618.

20. James Kirkpatrick, Razvan Pascanu, Neil C. Rabinowitz, et al. Overcoming catastrophic forgetting in neural networks. Proceedings of the National Academy of Sciences, 2017. doi: 10.1073/pnas.1611835114.

21. Yao Zhang, James Jordon, Ahmed M. Alaa, and Mihaela van der Schaar. Lifelong bayesian optimization. arXiv, 2019. doi: 10.48550/arXiv.1905.12280.

22. Zaixi Zhang, Ruofan Jin, Linlin Chao, Guangxue Xu, Yikun Zhang, Guowei Zhou, D. Yin, Yingqing Guo, Yaqi Fu, Yukang Yang, et al. Rnagenesis: a generalist foundation model for functional rna therapeutics. bioRxiv,pages 2024–12, 2024.

23. Leland McInnes, John Healy, and James Melville. Umap: Uniform manifold approximation and projection for dimension reduction. arXiv preprint arXiv:1802.03426, 2018.

